# Sex differences in responses to aggressive encounters among California mice

**DOI:** 10.1101/2023.12.12.571325

**Authors:** Jace X. Kuske, Alexandra Serna Godoy, Alison V. Ramirez, Brian C. Trainor

## Abstract

Despite how widespread female aggression is across the animal kingdom, there remains much unknown about its neuroendocrine mechanisms, especially in females that engage in aggression outside the peripartum period. Although the impact of experience in aggressive encounters on steroid hormone responses have been described, little is known about the impact of these experiences on female behavior or their neuropeptide responses. In this study, we compared behavioral responses in both male and female adult California mice based on if they had 0, 1, or 3 aggressive encounters using a modified resident intruder paradigm. We measured how arginine vasopressin and oxytocin cells in the paraventricular nucleus responded to aggression using c-fos immunohistochemistry. We saw that both sexes disengaged with intruders with repeated aggressive encounters, but that on the final day of testing females were most likely to freeze when they encountered intruders compared to no aggression controls – which was not significant in males. Finally, we saw that percent of arginine vasopressin and c-fos co-localizations in the posterior region of the paraventricular nucleus increased in males who fought compared to no aggression controls. No difference was observed in females. Overall, there is evidence that engaging in aggression induces stress responses in both sexes, and that females may be more sensitive to the effects of fighting.

## Introduction

Aggression has many uses and causes in animals, and many different species display aggression in a wide range of contexts. In some species females use aggression for the same reason as their male counterparts - in defense or acquisition of resources (Been et al., 2019; Duque-Wilckens & Trainor, 2017). This could be performing aggression in defense of a territory (Goymann et al., 2008; Rosvall, 2008), in defense of a mate (Bales & Carter, 2003; Bowler et al., 2002; Goymann et al., 2008), or to maintain dominance status (Lord et al., 2021). Many studies investigating the neuroendocrine mechanisms of aggression are conducted in rodents, and in the most commonly studied species females do not defend territories but will exhibit aggression to defend pups (Lonstein & Gammie, 2002) or mating opportunities (Aubry et al., 2022; Newman et al., 2019). Thus, there is strong evidence that females across a variety of species engage in aggressive behavior. An important question is how engaging in aggressive behavior impacts the neuroendocrine system.

An extensive literature has outlined how engaging in aggressive encounters alters brain and behavior in male vertebrates (Marler & Trainor, 2020). In male California mice (*Peromyscus californicus*), winning aggressive encounters triggers a transient increase in testosterone levels and more enduring increases in androgen receptor expression in the mesolimbic dopamine system (Fuxjager et al., 2009, 2010). These changes are thought to contribute to what is dubbed the winner-challenge effect, wherein animals’ increased aggression occurs following winning experiences thus allowing an individual to win future aggressive encounters (Oyegbile & Marler, 2005). Similar results have been reported in humans (Carré, 2009; Casto & Edwards, 2016).

Comparatively less is known about how winning experiences affect female brain and behavior. Female California mice demonstrated shorter attack latencies across three consecutive resident-intruder tests but showed no changes in other aggressive behaviors (Silva et al., 2010). While male California mice show higher testosterone levels after winning an encounter, females have lower plasma levels of progesterone (Davis & Marler, 2003) and higher levels of oxytocin (OXT) and corticosterone (Trainor et al., 2010) after winning aggressive encounters. These data show that there are important differences in how engaging in aggression affects males and females. The increased corticosterone finding in particular suggests that engaging in aggression could be aversive to females. Recent work in outbred CD-1 mice demonstrates that for some males engaging in aggressive behavior is reinforcing while for most females engaging in aggression is not reinforcing but they do not mention if it is aversive (Aubry et al., 2022).

Instead, though female CD-1s seek and prefer social interaction with other females (Ramsey et al., 2022). Thus although females can engage in aggression, this may be an aversive experience compared to simply interacting with another female. A key question is whether these effects are context dependent. Female *Mus musculus* do not defend territories, leaving the question of whether aggression would be aversive to females that naturally defend territories.

California mice are a unique species of deer mouse that display monogamy and perform joint territorial aggression (Rieger et al., 2021), and females engage in aggression regardless of if pups or mates are present (Davis & Marler, 2003; Duque-Wilckens & Trainor, 2017; Silva et al., 2010). Previous work has identified mixed relationships with progesterone, OXT, and arginine vasopressin (AVP) in female California mouse aggression (Davis & Marler, 2003; Steinman et al., 2015; Trainor et al., 2010). In these same studies, regions like the bed nucleus of the stria terminalis and the paraventricular nucleus (PVN) have also been implicated in California mouse aggression. However, little is known about the impacts of aggression on the brains and behavior of females compared to what we know about male California mouse aggression. The primary goals of this study are to identify how female residents responded to intruders, and to find how previously identified neuropeptide neurons OXT and AVP change in response to aggression within the PVN.

## Materials and methods

### Animals

All animals were bred in-house and were communally housed with same sex siblings prior to testing and housing manipulations. Each mouse was housed in cages of 3 to 4 with Sani-Chips bedding with cotton nestlets in clear polypropylene cage under a 16L:8D light cycle (lights on at 2PM) and *ad libitum* access to water and food (Harlan Teklad 2016). All procedures used were approved by the University of California, Davis Institutional Animal Care and Use Committee (IACUC).

### Territorial aggression

A total of 42 adult California mice were used (female = 24, male = 18) that were around 4 to 8 months of age at time of testing. All animals were tested across three consecutive days and randomly assigned into one of three groups: zero, one, or three aggressive encounters. All animals were single housed three days before the first day of testing.

All intruders were shaved on their rump to help identify them from the residents.

Territorial aggression was measured by a modified version of the classic resident-intruder test. Briefly, an intruder was placed in the opposite corner of the resident’s cage and allowed to directly interact with the resident for a total of 10 minutes. Two tests were ended early to ensure safety of intruders and prevent injury during aggressive encounters. Aggression and in-cage anxiety behaviors were video recorded and hand scored using BORIS by observers blind to treatment groups (Table 1). One hour after the end of the final aggression test on day 3, each animal was euthanized using a transcardial perfusion using 4% PFA in PBS. Brains were stored in PFA overnight and then switched to 30% sucrose in PBS for 2-3 days, then they were flash frozen with dry ice and stored in -80 freezer until sections were cut on a cryostat.

**Table 1.** List of different behaviors measured on BORIS. Frequency, duration, and latency to perform each behavior was recorded and included in statistical analyses.

| Behavior | Description |
| --- | --- |
| **Aggression** |  |
| Chasing | Resident pursuing/running after intruders |
| Wrestling | Physical contact with intruders, accompanied by tumbling together with resident likely biting the intruder (though not always clearly visible) |
| Anogenital sniffing | Resident sniffing the anogenital region of the intruder |
| Boxing | Resident defensively uses forepaws to swat at and lunge towards intruder |
| **Anxiety** |  |
| Freezing | Resident is immobile and no part of its body is moving for at least 2 seconds, often visibly tense |
| Autogrooming | Resident begins grooming itself, most often after wrestling bout or any form of physical contact with intruder. |

### Immunohistochemistry procedure

To identify how the brain changes in response to encounters with intruders, we stained for OXT, AVP, and c-Fos expression in the anterior and posterior paraventricular nucleus (aPVN, pPVN) using a three day triple label protocol. After the brains were frozen, they were cut at 40 μm sections. Sections were chosen starting from -0.36 bregma (BrainMaps.org) and selecting one every 3-4 slices from the anterior to posterior of the PVN. Sections were washed twice for 5 minutes in PBS and incubated in a 10% normal goat serum (NGS) in PBS blocking solution for one hour and washed once more. Then, sections were incubated overnight in a rabbit anti-cFos antibody (226 008, 1:1000, Synaptic Systems, Göttingen, Germany) in 2% NGS and PBS with Triton-X (PBS-TX) at room temperature. On day 2, sections were washed three times in PBS and incubated for 2 hours in a goat anti-rabbit Alexa Fluor 555 antibody (A21429, 1:500, Invitrogen, Carlsbad, CA, USA), they were washed in PBS three more times, and then were incubated overnight in 4° in a cocktail of mouse anti-OXT (MAB5296, 1:1000, EMD Millipore Corp, Darmstadt, Germany) and guinea pig anti-AVP (403 004, 1:1000, Synaptic Systems, Göttingen, Germany). On day 3, tissue was washed three times in PBS and then incubated for an hour at room temperature in a secondary cocktail with goat anti-mouse Alexa Fluor 488 (A11001, 1:500, Invitrogen, Carlsbad, CA, USA) and biotinylated goat anti-guinea pig (BA- 7000, 1:500, Vector Laboratories, Burlingame, CA, USA), followed by three more PBS washes and a 30 minute incubation in streptavidin conjugated to Alexa Fluor 405 (S32351, 1:500, Invitrogen, Carlsbad, CA, USA) diluted in NGS and PBS-TX, and three final PBS washes. Slices were mounted onto Superfrost plus slides (Fisher, Pittsburgh, PA, USA), left to dry, and then coverslipped with Vectashield (Vector Laboratories, Burlingame, CA, USA).

### Immunohistochemistry analysis

Images were taken using a monochromatic Axiocam MRm camera (Zeiss Meditec) connected to a Ziss Axioimager. To quantify the number of cells in each region and their co- localizations, we used ImageJ to create boxes (0.26 x 0.35 mm) for areas of interest and record immunoreactivity in OXT, AVP, and c-fos cells. Quantification was done by a researcher blind to sex and conditions of each animal.

### Statistical analyses

All statistical analyses were performed using RStudio. For measures of aggression (chasing, wrestling) measures of anxiety (freezing) and for other social behaviors (anogenital sniffing), we used repeated measures ANOVA with pairwise *t*-test as a post hoc test. Because of heterogeneous variance across groups, we used non-parametric for both behavior and cell counting data. Specifically, we used the Kruskal-Wallis rank sum test for comparisons of aggression, anxiety and other social behaviors between our aggression groups, and for comparisons of cell number and percent co-localizations of AVP, OXT, and c-fos in the PVN.

## Results

### Multiple territorial intrusions reduces engagement with intruders

For mice that were tested in 3 aggression tests, we observed unexpected changes in social investigation and aggression. Male and female residents sniffed the anogenital region of the intruder mice less after day 1 (Fig. 1B; F(1, 26) = 22.69, p = 6.27e-05). This decreased engagement was associated with a decrease in the frequency of wrestling by the final day of testing (Fig. 1D; F(1, 27) = 4.803, p = 0.037). Overall, females engaged in overt displays of aggression less frequently than males with lower rates of chasing (Fig. 1C; F(1,12) = 9.072, p = 0.011) and wrestling (Fig 1D; F(1,12) = 8.044, p = 0.015). Females also showed a higher latency to wrestle compared to males (Fig. 1E; F(1, 12) = 11.28, p = 0.006).

**Figure 1.**
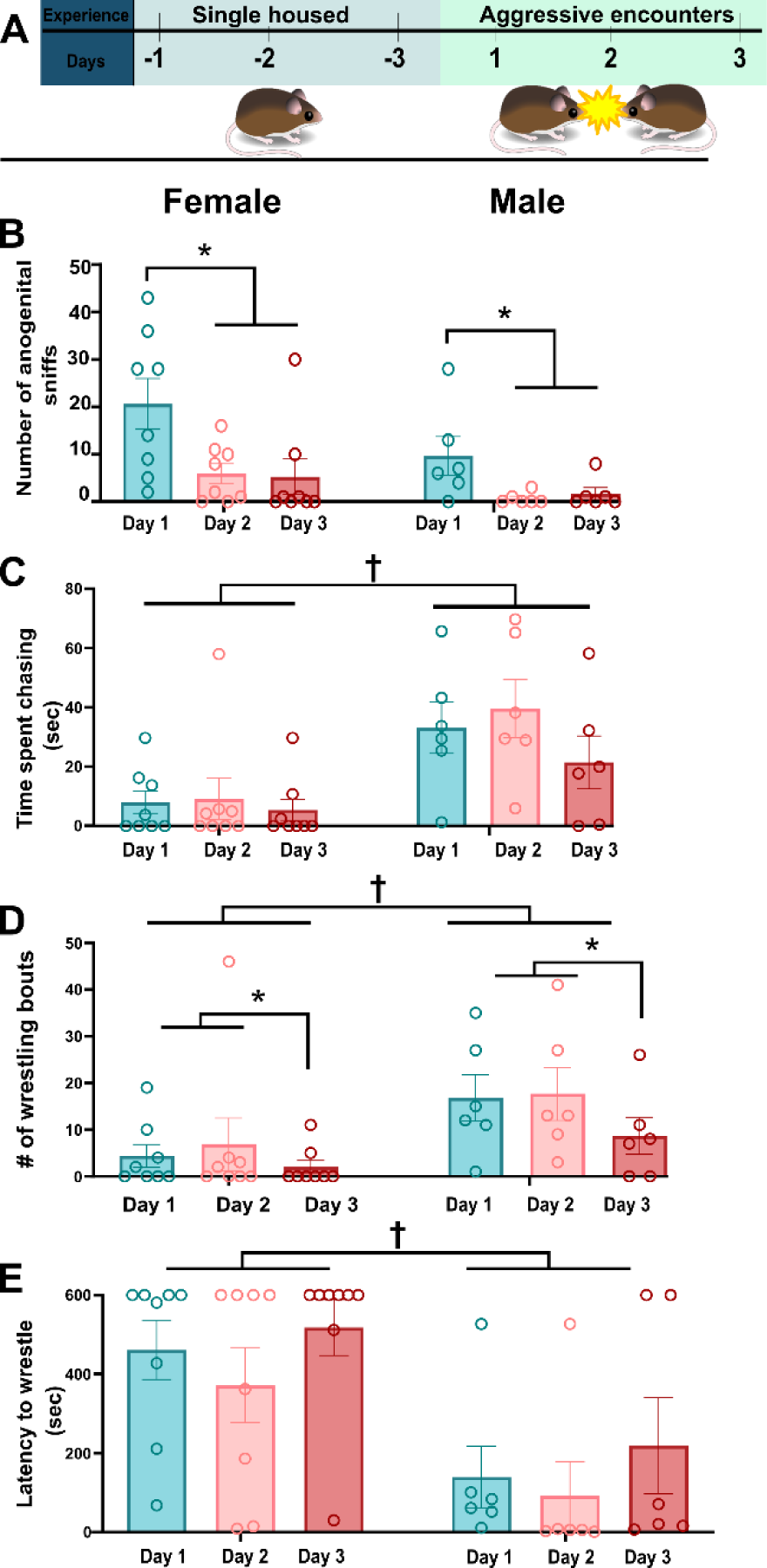
Timeline and changes of aggressive behaviors across repeated encounters. * indicate significant changes in behavior across day, † indicate significant sex differences. A) Testing and housing manipulation timeline. B) For both sexes, repeated aggressive encounters reduced frequency of anogenital sniffing (p = 6.27e-05). C) Overall, males chased the intruders more often than females did, but this did not change across repeated tests (p = 0.0108). D) There was a decrease in wrestling bouts by the final day of tests (p = 0.0393) with male residents engaging in wrestling with intruders more often than female residents (p = 0.015). E) There was a significant difference in latency to wrestle with intruders with female residents taking longer to wrestle compared to male residents (p = 0.006).

### Social investigation behaviors in resident intruder test

On the final day of testing mice assigned to three encounters showed less anogenital sniffing (Fig. 2B, Kruskal-Wallis H(1) = 7.8163, p = 0.005), and took longer to sniff (Fig. 2C; H(1) = 6.19, p = 0.01) compared to mice assigned to one encounter. Decreased social investigation might reflect an increase in aggression or an increase in anxiety-related behavior. However, we observed was no change in frequency of chasing (Fig. 2D; H(1) = 0.35, p = 0.56) nor wrestling bouts (H(1) = 2.12, p = 0.15). There were no differences in boxing or nose-nose sniffing (Table 2).

**Figure 2.**
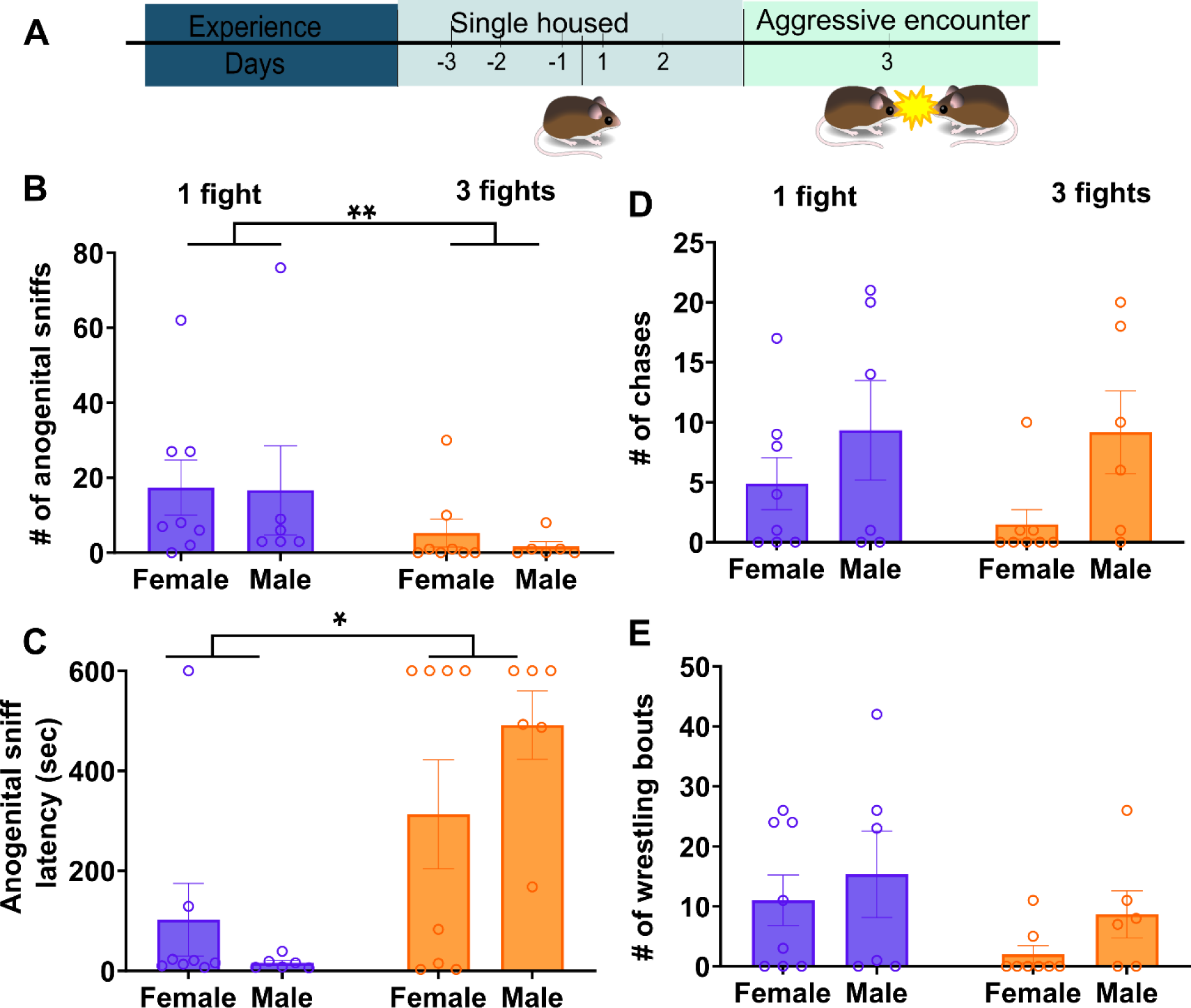
Changes in social behaviors based on aggressive experience. ** indicates significant than less than 0.001 level, * is 0.01. B) Timeline of animals with a single aggressive encounter, “no aggression” group was single housed for duration of the experiment. B) Repeated fights resulted in decreased engagement of intruders through decrease in frequency of anogenital sniffs (H(1) = 7.82, p = 0.005), C) Repeated fights resulted in greater latency to engage with intruders (H(1) = 6.19, p = 0.01). D) There was no change in overall aggression with chasing nor (E) wrestling changing as a result of repeated encounters.

**Table 2.** Mean ± SE of various behavior on final day of testing by group and sex.

| **Behavior** | **No**<br>**aggression**<br>**female** | **No**<br>**aggression**<br>**male** | **1 fight**<br>**female** | **1 fight**<br>**male** | **3 fights**<br>**female** | **3 fights**<br>**male** |
| --- | --- | --- | --- | --- | --- | --- |
| **Boxing** | - | - | 1.3 ±<br>0.61 | 0.5 ± 0.38 | 0 ± 0 | 1.83 ±<br>1.45 |
| **Nose-nose**<br>**sniffing** | - | - | 9 ±<br>2.17 | 3.75 ±<br>1.76 | 13.67 ±<br>7.74 | 4.83 ±<br>2.29 |
| **Freezing** | 0.5 ± 0.27 | 3.0 ± 1.61 | 7.25 ±<br>2.69 | 6.83 ±<br>3.33 | 8.13 ±<br>2.41 | 10.3 ±<br>2.79 |
| **Autogrooming** | 1.75 ±<br>0.53 | 3.5 ± 1.60 | 4.25 ±<br>1.04 | 2.67 ±<br>0.67 | 2.13 ±<br>0.79 | 3.33 ±<br>1.17 |

### Anxiety-like behaviors present in resident intruder test

Female residents exposed to an intruder showed passive stress coping behavior like freezing when confronted (Fig. 3A; H(1) = 9.65, p = 0.002) whereas male residents showed no significant change in freezing (H(1) = 2.03, p = 0.15). Intriguingly, there is no difference in active stress coping behavior like autogrooming in either sex or condition (H(1) = 1.11, p = 0.29).

**Figure 3.**
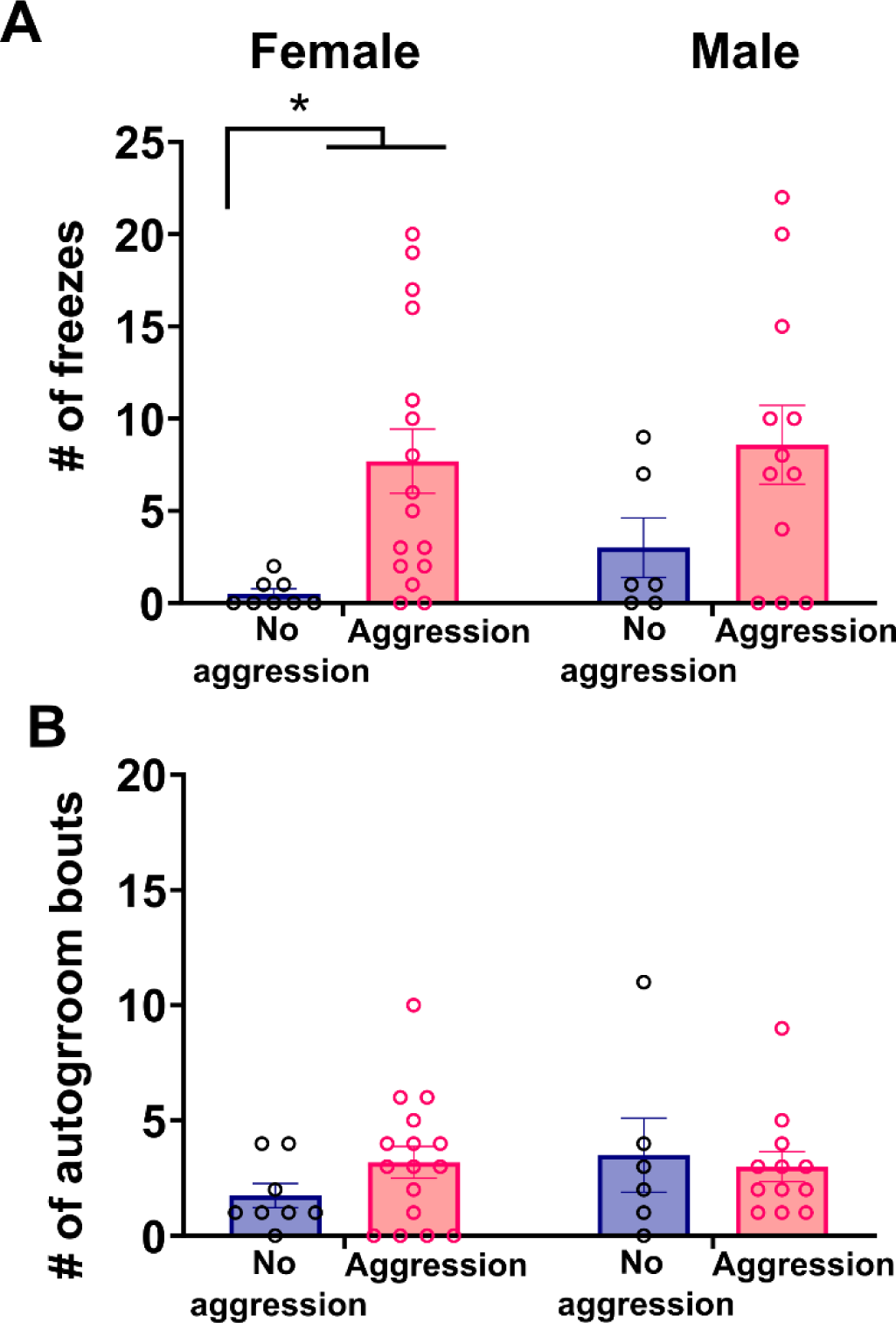
Changes in anxiety-like behaviors based on aggressive experience. ** indicates significant than less than 0.01 level. A) Female residents froze in response to intruders only (H(1) = 9.65, p = 0.002), but males showed no difference in when they froze. C) There were no sex or group differences in number of autogrooming bouts.

### Neuroendocrine responses to intruder encounters

AVP/c-fos co-localizations were observed in posterior and anterior PVN (Fig. 4). In the posterior PVN, males (Fig. 5A; H(1) = 9.59, p = 0.002) but not females with aggressive experience (combining “1 fight” and “3 fights” conditions) had more percent AVP/c-fos co- localizations compared to residents with no exposure to intruders (no aggression condition). This was in conjunction with no changes in total number of AVP cells in the posterior PVN (Fig. 5B; H(1) = 1.20, p = 0.27), indicating that changes in co-localizations were not driven by changes in AVP cell number. In the anterior PVN, there were no differences in AVP/c-fos co-localizations but females exposed to 1 or 3 encounters had fewer total AVP cells than females in the no aggression group (Fig. 5C; H(1) = 5.30, p = 0.02). No differences in AVP cell counts were observed in males. For OXT, effects of aggressive encounters on OXT/c-fos co-localizations in the posterior PVN were less robust, as there was no significant differences in males or females. However, when males and females were pooled together, the no encounter group had fewer OXT/c-fos co-localizations compared to animals that were exposed to an intruder (Fig. 3, H(1) = 5.09, p = 0.02). However, no other differences in co-localizations or cell counts were found in either the anterior or posterior region of the PVN (Table 3).

**Figure 4.**
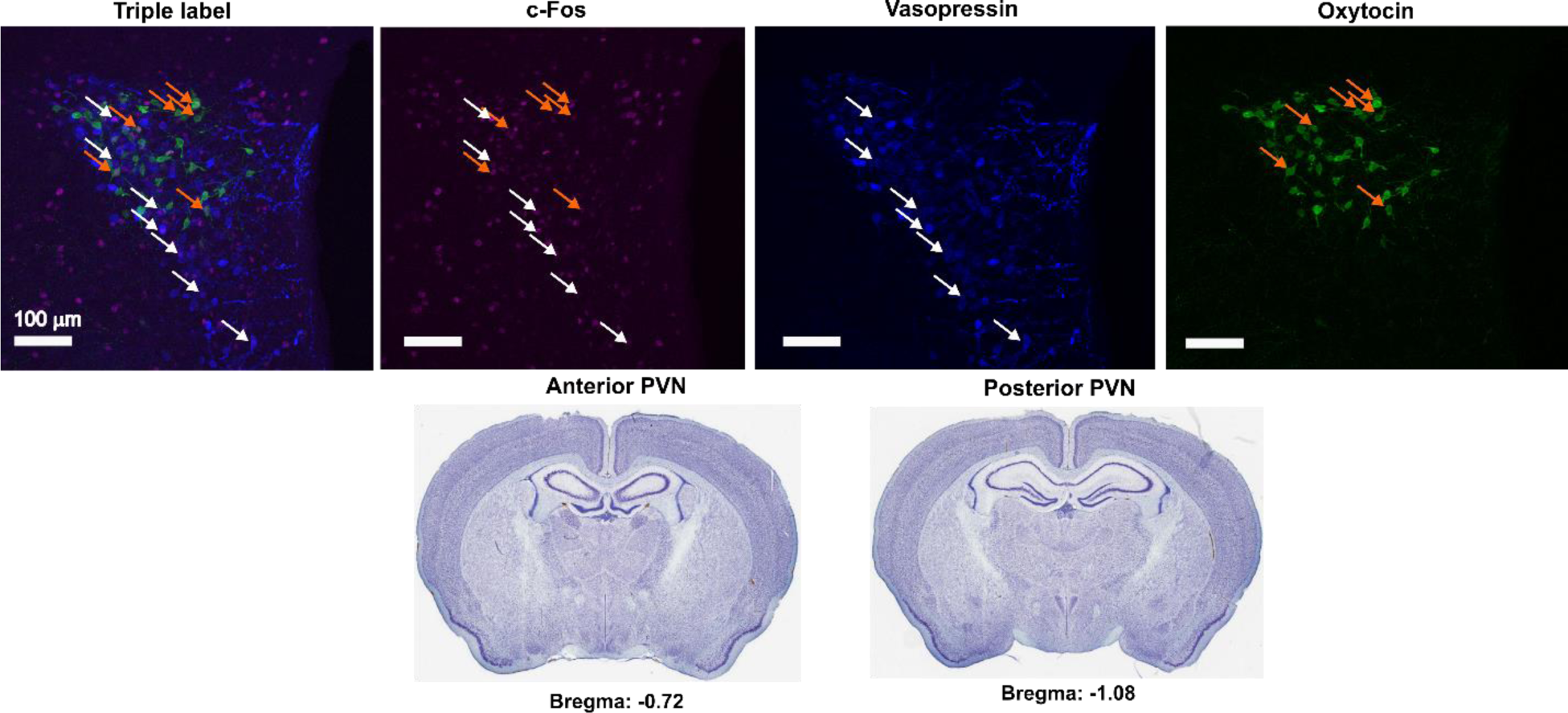
Image of positive staining in the posterior PVN. White arrows indicate AVP/c-fos co- localizations, orange arrows indicate OXT/c-fos co-localizations. Below is shown relatively where we labeled the anterior and posterior PVN.

**Figure 5.**
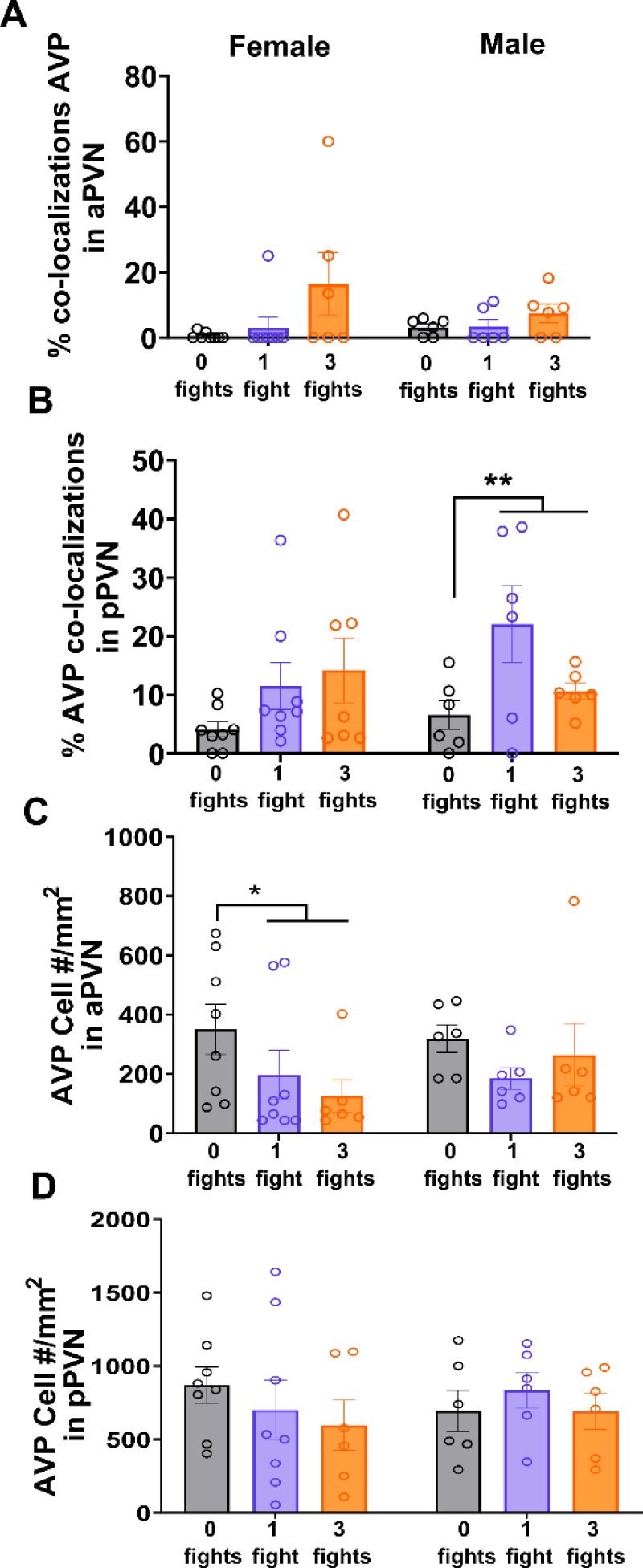
Changes in vasopressin following aggressive encounters. A) No differences in group or sex in the percent AVP/c-fos co-localizations (n.s.). B) Males exposed to intruders had greater percent AVP/c-fos co- localizations compared to no aggression controls (H(1) = 9.59, p = 0.002). C) Interestingly, no aggression controls had greater amounts of cells than those exposed to intruders (H(1) = 5.30, p = 0.02. D) There were not greater amounts of AVP cells in the posterior region (n.s.).

**Figure 6.**
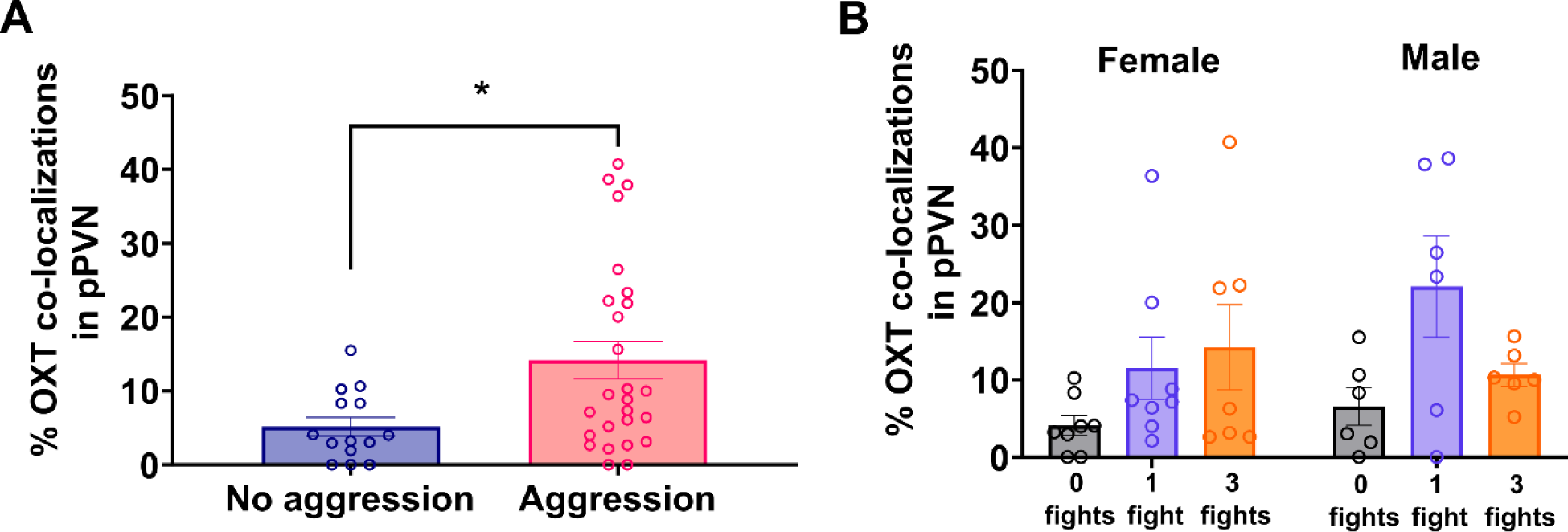
Changes in percent OXT/c-fos co-localizations based on aggressive experience. ** indicates significant than less than 0.01 level. A) There is greater percent OXT/c-fos co- localizations on in those with aggressive experience compared to those without experience - when sexes are combined (H(1) = 5.01, p = 0.02) in the pPVN. B) This difference does not appear when separating by sex.

**Table 3.** Mean SE of total cell counts of OXT in two different regions of the PVN.

| **Region and**<br>**count** | **No**<br>**aggression**<br>**female** | **No**<br>**aggression**<br>**male** | **1 fight**<br>**female** | **1 fight**<br>**male** | **3 fights**<br>**female** | **3 fights**<br>**male** |
| --- | --- | --- | --- | --- | --- | --- |
| **OXT aPVN** | 59.3 ± 12.5 | 60 ± 17.8 | 39 ± 4.8 | 60 ± 12.2 | 53.1 ± 11.6 | 46.7 ± 13.1 |
| **OXT pPVN** | 56 ± 7.2 | 61 ± 8.7 | 56.6 ± 11.3 | 44.2 ± 7.7 | 36.6 ± 9.9 | 46.3 ± 11.2 |

## Discussion

Overall, repeated encounters with intruders decreases the amount of aggression and engagement with intruders displayed by male and female residents. Additionally, we observed that females but not males show passive stress coping behavior in response to intruders.

Aggressive encounters had different effects on AVP immunoreactivity in females and males. In females, engaging in aggressive encounters reduced the number of AVP immunoreactive cells in the anterior region of the PVN without affecting c-fos/AVP co-localizations. In contrast, males exposed to intruders had higher percent AVP/c-fos co-localizations in the posterior region of the PVN but did not alter the number of immunoreactive cells. Together, these data indicate that engaging in aggressive encounters has different behavioral and neurobiological consequences in females and males.

### Aggressive encounters induce stress coping behavior in females

Although some studies have reported that engaging in aggression can be reinforcing, other work has highlighted that engaging in aggression can be an aversive experience (Golden et al., 2016). Previous work in female California mice showed that engaging in aggression triggers a delayed decrease in progesterone as opposed to increased testosterone reported for males (Davis & Marler, 2003). Although one study reports that females show reduced attack latencies with repeated aggressive encounters (Silva et al., 2010), they also showed no changes in total number of offensive attacks. Other results showed that females that engaged in aggression triggered an acute increase in corticosterone compared to females that were not exposed to intruders (Trainor et al., 2010). Our finding that repeated encounters with intruders decreased overall engagement with intruders and total aggressive fights suggests that engaging in aggression may be aversive for female California mice. Indeed, losing aggressive encounters also has more enduring effects on social approach behaviors in females than in males (Greenberg et al., 2014; Trainor et al., 2011). A surprising finding was that engaging in repeated aggressive encounters did not increase aggressive behavior in males, as reported in previous studies (Oyegbile & Marler, 2005; Trainor & Marler, 2001). A key difference in previous work was that males were paired with a female whereas in our work all mice were sexually inexperienced and singly housed for 3 days prior to testing. California mice form pair bonds (Gubernick, 1988; Rieger et al., 2019), which have important effects on stress responses (Bales et al., 2021). Our results suggest that pair bonding may alter the valence of aggressive interactions, especially for males. Indeed in this species, pairs become more coordinated and behaviorally “synch up” the longer they cohabitate (Rieger et al., 2021) - indicating that having a partner is a significant feature in the natural aggressive experience of these mice. Similarly, female California mice avoided aggressive encounters when they were alone, but approached intruders more if a mate was present (Rieger et al., 2021). Another factor that may contribute to the difference in results is the lack of training. In previous studies with males, they were trained to attack younger, smaller, mildly anesthetized intruders, whereas our mice fought against younger or evenly matched intruders that were not anesthetized. Overall, pairing status, previous experience, and housing conditions can all impact the behavioral effects of aggressive experiences.

### Neuroendocrine changes following aggressive encounters

In male California mice, aggressive experience increased AVP/c-fos co-localizations whereas in females aggressive experienced reduce AVP cell counts. There are important sex differences in how AVP regulates aggression. For example there was a positive relationship between biting and the number of positive AVP cells in the bed nucleus of the stria terminalis in male California mice but not females (Steinman et al., 2015). In female Syrian hamsters housed in long photoperiods, AVP in the anterior hypothalamus inhibits aggression (Gutzler et al., 2010) in contrast to males housed in long photoperiods where AVP increased aggression (Caldwell & Albers, 2004). In sexually inexperienced female rats, microdialysis showed reduced AVP release in the dorsal lateral septum and that V1a receptors were required for the expression of social aggression (de Moura Oliveira et al., 2021). In male rats, AVP release in the lateral septum during aggression is strain dependent, with more aggressive rats showing decreased AVP release and less aggressive rats showing increased AVP release (Beiderbeck et al., 2007). Although V1aR antagonist infusion did not affect aggressive behavior in these lines, V1aR were found to be critical for facilitating experience-dependent increases in aggression in sexually experienced male rats (Veenema et al., 2010). Interestingly deletion of AVP cells in the PVN increased social investigation in female mice but not in males (Rigney et al., 2021).

Sex differences in the behavioral effects of AVP have been observed in other contexts.

For example, AVP1a antagonists infused into the lateral septum increase social play in male rats but decrease play in females (Bredewold et al., 2023). Similarly social defeat in California mice has stronger effects on AVP mRNA and immunoreactivity in the PVN in males than females, and interestingly males showed reduced AVP positive cells after defeat (Steinman et al., 2015). This seems specific to social stress as non-social stressors like chronic variable stress in male California mice produced greater AVP mRNA in the PVN, though females were not studied here (De Jong et al., 2013). We observed that only males with aggressive experience showed increased percent AVP/c-Fos co-localizations compared to those with no experience. Increases in co-localizations suggest an increase in neural activity and possibly increased AVP release. In females that experienced aggression, there was a decrease in AVP positive cells in the PVN. A decrease in immunoreactivity could indicate a decrease in AVP synthesis or an increase in AVP release. Further study is needed to distinguish between these two possible outcomes.

Across males and females, we observed increased OXT/c-fos co-localizations in animals that engaged with intruders compared no aggression controls. Interestingly, OXT neurons in the PVN are responsive to social stress, which corresponds with our observations that engaging in aggressive encounters enhanced stress-related behaviors. In California mice that experience social defeat, there are greater OXT/c-fos co-localizations in the PVN (Steinman et al., 2016).

Interestingly, a previous study showed that following resident intruder tests female but not male California mice showed greater OXT positive cells in the PVN during long photoperiods (Trainor et al., 2010). This might mean that both sexes are susceptible to the stress of losing fights, but that females might be more sensitive to any fighting.

## Conclusions

Overall, our results suggest that engaging in aggression elicits behavioral stress responses in both female and male California mice that are sexually inexperienced. Further work is needed to assess the impact of pairing status on stress responses elicited during aggressive behavior.

While paired male California mice show increased aggression with repeated experience, how aggressive experience impacts female aggression has never been examined. There is a great opportunity to explore how pairing status may influence aggressive behavior and how the brain responses to this change. Pair bonding has important effect on OXT and AVP signaling pathways, which in turn will likely modulate the impact of pairing status on aggression.

## Acknowledgements

This work supported by the Eugene Cota-Robles Fellowship and the NSF Graduate Research Fellowship to JXK and NSF IOS 1937335 to BCT. Thanks to A. Baxter, A. Seelke, J. Bond, and K. Bales for helpful discussions.

